# Loss of Kat2A Enhances Transcriptional Noise and Depletes Acute Myeloid Leukemia Stem-Like Cells

**DOI:** 10.1101/446096

**Authors:** Ana Filipa Domingues, Rashmi Kulkarni, George Giotopoulos, Shikha Gupta, Shengjiang Tan, Elena Foerner, Rita Romano Adao, Keti Zeka, Brian J. Huntly, Sudhakaran Prabakaran, Cristina Pina

**Author notes:** These authors contributed equally to this work (Ana Filipa Domingues, Rashmi Kulkarni).

## Abstract

Acute Myeloid Leukemia (AML) is an aggressive hematological malignancy with abnormal progenitor self-renewal and defective myelo-monocytic differentiation. Its pathogenesis comprises subversion of transcriptional regulation, through mutation and by hijacking normal chromatin regulation. Kat2a is a histone acetyltransferase central to promoter activity that we recently associated with stability of pluripotency networks, and identified as a genetic vulnerability in AML. Through combined chromatin profiling and single-cell transcriptomics, we demonstrate that Kat2a contributes to leukemia propagation through homogeneity of transcriptional programs and preservation of leukemia stem-like cells. Kat2a loss reduces transcriptional bursting frequency in a subset of gene promoters, generating enhanced variability of transcript levels but minimal effects on mean gene expression. Destabilization of target programs shifts cellular equilibrium out of self-renewal towards differentiation. We propose that control of transcriptional variability is central to leukemia stem-like cell propagation, and establish a paradigm exploitable in different tumors and at distinct stages of cancer evolution.

## INTRODUCTION

Acute Myeloid Leukemia (AML) is the most prevalent leukemia in adults with a dismal prognosis of less than 30% 5-year survival (Dohner et al., 2017). It is a heterogeneous disease, clinically and pathologically, with common cellular themes of myeloid differentiation block, and recurrent molecular targeting of chromatin and transcriptional regulation. Effects on transcription are reflected in the AML mutational spectrum (Cancer Genome Atlas Research et al., 2013), as well as through the implication of general transcriptional co-regulators in AML pathogenesis, in the absence of specific mutation events (Roe and Vakoc, 2013). Examples of these are specific AML dependencies on BRD4 (Dawson et al., 2011; Zuber et al., 2011), LSD1 (Harris et al., 2012) or DOT1L (Bernt et al., 2011; Daigle et al., 2011). Moreover, chemical inhibitors exist to target these regulators and have progressed to clinical trials (Gallipoli et al., 2015). More recently, TFIID and SAGA subunit TAF12 was shown to be critical for MYB protein stability and transcriptional activity in AML cells through its participation in the TFIID complex (Xu et al., 2018).

In a recent CRISPR drop-out screen of genetic dependencies in AML, we have identified several members of the SAGA complex, including histone acetyl-transferase *KAT2A*, as being required for AML maintenance (Tzelepis et al., 2016). KAT2A was suggested to impact cell survival and differentiation status, but its precise molecular mechanisms of action remain to be elucidated, and it is unclear whether it is required in AML initiation, as well as maintenance. *Kat2a* is a mammalian orthologue of yeast histone acetyl-transferase *Gcn5*, and is required for H3K9 acetylation (H3K9ac) (Jin et al., 2014), a modification that fine-tunes, rather than initiates, locus-specific transcriptional activity. Kat2a is required for specification of mesodermal derivatives during early embryonic development (Lin et al., 2007) (Wang et al., 2018), and for survival of neural stem and progenitor cells (Martinez-Cerdeno et al., 2012). Loss of *Kat2a* has minimal impact on normal hematopoiesis, except for promotion of terminal granulocyte differentiation through release of protein acetylation-dependent inactivation of Cebpa transcriptional activity (Bararia et al., 2016).

Yeast Gcn5 is a classical regulator of intrinsic transcriptional noise (Raser and O'Shea, 2004), with deletion mutants enhancing cell-to-cell variability in gene expression measured across a range of locus fluorescence reporters (Weinberger et al., 2012). Transcriptional noise reflects the variability in the number of mRNA molecules produced from a given locus through time; with snapshot studies of gene expression capturing the same phenomenon as cell-to-cell transcriptional heterogeneity (Sanchez et al., 2013). Transcriptional noise is consequent to the bursting nature of gene expression (Chubb and Liverpool, 2010): for most if not all loci, transcriptional activity is not continuous, but burst-like or episodic, with locus-specific rates of locus activation (**κ**ON), inactivation (**κ**OFF), and RNA production (**κ**RNA), as well as RNA degradation (Raj et al., 2006) contributing to the net effect. Frequency of bursting depends on the **κ**ON rate, whilst **κ**RNA impacts the burst size (Cai et al., 2006). Both parameters contribute to mean gene expression, whilst transcriptional noise is more strictly dependent and anti-correlated with burst frequency (Hornung et al., 2012). In yeast, size and frequency of bursts are influenced by histone acetylation in gene bodies and promoters, respectively (Weinberger et al., 2012).

In functional terms, transcriptional noise has been directly implicated as a mechanism of cell fate choice in yeast (Blake et al., 2006) and bacteria (Suel et al., 2006), and recurrently associated, albeit correlatively, with cell fate transitions in mammalian systems (Moris et al., 2016). We had previously shown that normal transitions into hematopoietic lineage specification associate with cell-to-cell heterogeneity in gene expression (Pina et al., 2012; Teles et al., 2013). More recently, we have inhibited the activity of Kat2a in mouse embryonic stem cells, and observed an increase in transcriptional heterogeneity that impacted the stability of pluripotency with reconfiguration of correlation gene regulatory networks (GRNs) (Moris et al., 2018). Whilst we have not mechanistically linked enhanced heterogeneity with the loss of pluripotency, we observed propagation of variability of transcriptional levels through the GRNs downstream of nodes with differential H3K9ac. Cancer, and in particular leukemia, can be perceived as an imbalance between self-renewal and differentiation in favor of self-renewal. We postulated that enhancing transcriptional variability in AML cells would enhance the probability of cell fate transitions out of self-renewal into differentiation, with loss of leukemia stem-like cells (LSC). By using a retroviral-delivered *MLL-AF9* model of AML in a conditional *Kat2a* knockout background, we show that loss of *Kat2a* delays initiation of AML with locus-specific loss of H3K9ac and increased transcriptional variability amongst Kat2a histone acetylation targets. Modelling of transcriptional parameters suggests that the variability observed can be explained by changes in the frequency of bursting and thus reflects transcriptional noise. Cell clusters with transcriptional characteristics of stem cells are particularly affected, and we show that leukemia stem-like cells (LSC) are dramatic depleted in *Kat2a* knockout AML. Our data suggest that *Kat2a* contributes to the establishment and/or maintenance of LSC through buffering of gene expression variability. Furthermore, our results advance the notion that enhancing transcriptional noise can have pro-differentiation therapeutic potential in AML as well as other malignancies.

## RESULTS

### Conditional loss of Kat2a does not affect normal hematopoiesis and allows MLL-AF9-driven transformation

We sought to investigate *Kat2a* requirements *in vivo* during early leukemia initiation and progression, by generating a conditional *Kat2a^FI/FI*^ Mx1-Cre* mouse model and transforming *Kat2a* excised (KO) and non-excised (WT) bone marrow (BM) cells through retroviral delivery of an *MLL-AF9* fusion transcript. This strategy allows cellular and molecular investigation of *Kat2a* requirements during transformation, and critically, minimizes the putative confounding effect of acquired mutations on transcriptional heterogeneity, such as might be expected in established human or mouse AML. The choice of a strong oncogenic event such as *MLL-AF9*(Cancer Genome Atlas Research et al., 2013; Krivtsov et al., 2006; Somervaille and Cleary, 2006) also minimizes the need for cooperating genetic alterations. Specifically, we crossed *Kat2a^Fl/Fl^*C57Bl/6 mice (Lin et al., 2008) on a *Mx1-Cre^+/−^* background, and generated a stable mouse line homozygous for the *Flox* allele. *Mx1-Cre*-positive (KO) and *Mx1-Cre*-negative (WT) mice were compared across all experiments (Fig. 1A). We obtained locus excision by treatment of experimental and control mice with a course of intra-peritoneal polyinosylic-polycytidylic (*pIpC*) acid injections, as described (Chan et al., 2011). Excision was tested 4-6 weeks after injection and consistently achieved values greater than 80% in stem and progenitor cell compartments (Fig. 1B), reflected in a profound loss of gene expression, including amongst myeloid-biased (LMPP) and committed (GMP) progenitors critical for AML initiation (Goardon et al., 2011) (Fig. 1C). Of note, locus excision generates an in-frame product that joins the first 2 and the last exons (Supplementary Fig. 1A); this product is transcribed (Supplementary Fig. 1B), but should not code for catalytic or acetyl-binding activity (Supplementary Fig. 1A). In agreement with a previous report (Bararia et al., 2016), Kat2a was dispensable for HSC maintenance and function, as assessed by BM composition acutely after excision and throughout aging (Supplementary Fig. 1C-D), *in vitro* progenitor assays (Supplementary Fig. 1E), and by long-term reconstitution of irradiated recipients through transplantation (Supplementary Fig. 1F). Transformation of progenitor-enriched, lineage-depleted (Lin-) cells with an *MLL-AF9* fusion transcript was initially assessed *in vitro* through serial re-plating of WT and KO cells in in semi-solid medium-based colony-forming assays. Transformation was observed for cells of both genotypes, with similar efficiency (Fig. 1D). Locus excision (Fig. 1E) and gene expression loss (Fig. 1F) were maintained or even increased during transformation, suggesting that loss of Kat2a does not impede the initial selection of a leukemia-transformed clone.

**Figure 1.**
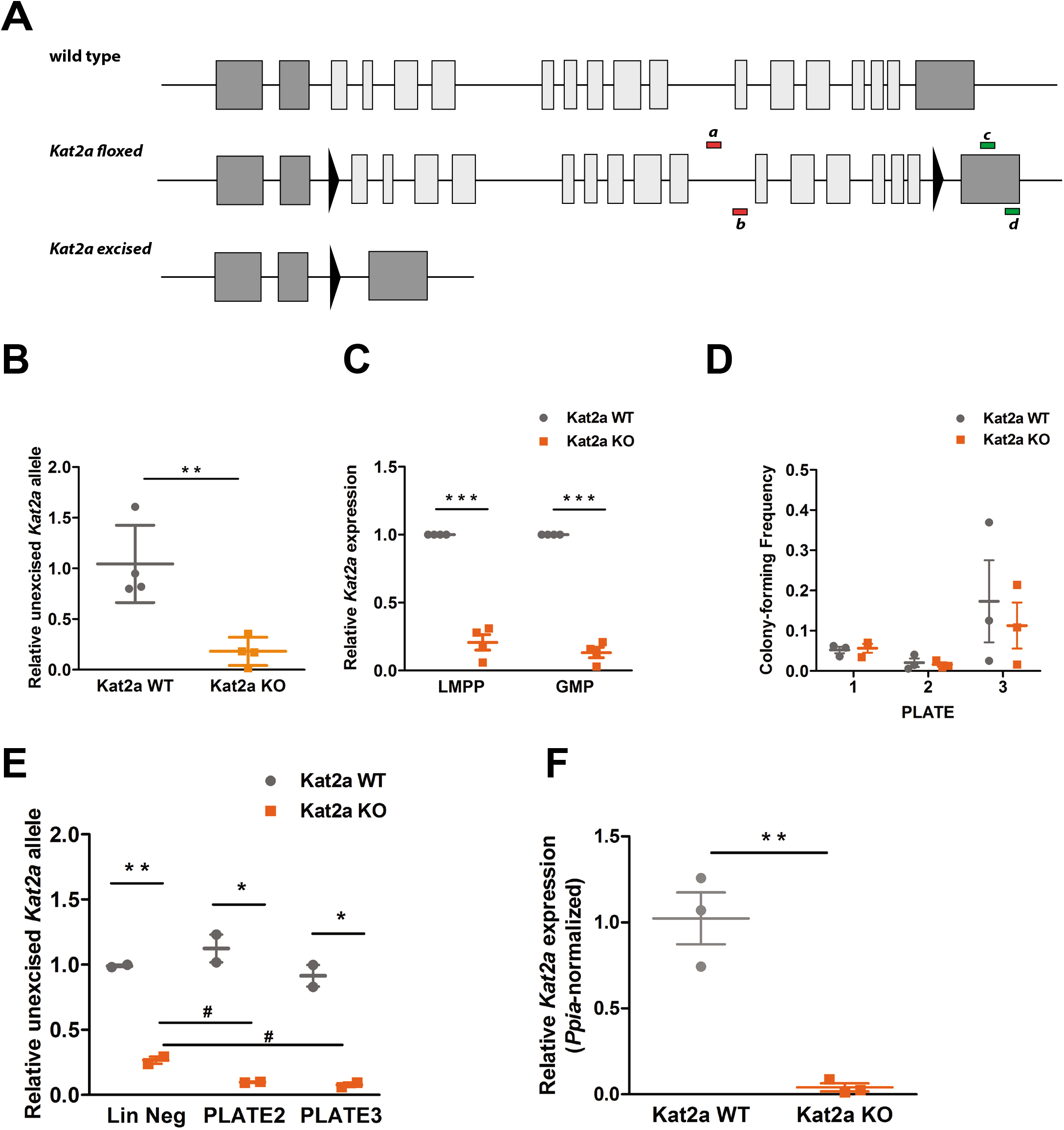
Conditional knockout of *Kat2a* is compatible with MLL-AF9-driven transformation. **(A)** Diagram of wild type *Kat2a* locus, conditional *Kat2a* floxed and the null Kat2a-excised allele, with excision detection strategy represented: *Kat2a* IN primers (within excised region)-*a* and *b*, and *Kat2a* OUT primers (downstream the excised region) - *c* and *d*. **(B)** Excision efficiency quantified by qPCR in mouse BM samples, mean ± SEM, n=4, ** p<0.01. **(C)** Quantification of *Kat2a* transcription levels in BM progenitor population, LMPP and GMP, by RT-qPCR mean ± SEM, n=2-5, ** p<0.001]. **(D)** Serial replating of Colony-forming cell (CFC) assays of MLL-AF9 transformed cells, mean ± SEM, n=3. **(E)** Excision efficiency was evaluated by qPCR during replating of MLL-AF9 transformed cells, mean ± SEM, n= 2-3, * p<0.01 and ** p<0.001. **(F)** Relative expression of *Kat2a* in MLL-AF9 primary leukemia BM cells from WT and KO backgrounds by RT-qPCR, mean ± SEM, n=3, ** p<0.001. Two-tailed t test was performed in (B), (C), (D), (E) and (F).

### Kat2a depletion impairs establishment of MLL-AF9-initiated leukemia

We investigated the longer-term effects of *Kat2a* loss in transformation progression *in vitro* by continued serial re-plating. Whilst no differences were seen in re-plating ability (Fig. 2A), it was noted that the colonies obtained had a clear component of differentiated cells (‘dubbed’ mixed or type II colonies (Johnson et al., 2003)) (Fig. 2B), which could also be observed in colonies initiated from primary BM transformed colonies (Fig. 2C). Accordingly, *Kat2a* KO colonies showed increased levels of the differentiation marker CD11b (Supplementary Fig. 2A). Interestingly, establishment of clonal liquid cultures from *in vitro* transformed cells revealed a relative advantage in culture initiation from WT cells (Fig. 2D), suggesting that an imbalance between self-renewal and differentiation in the KO setting.

**Figure 2.**
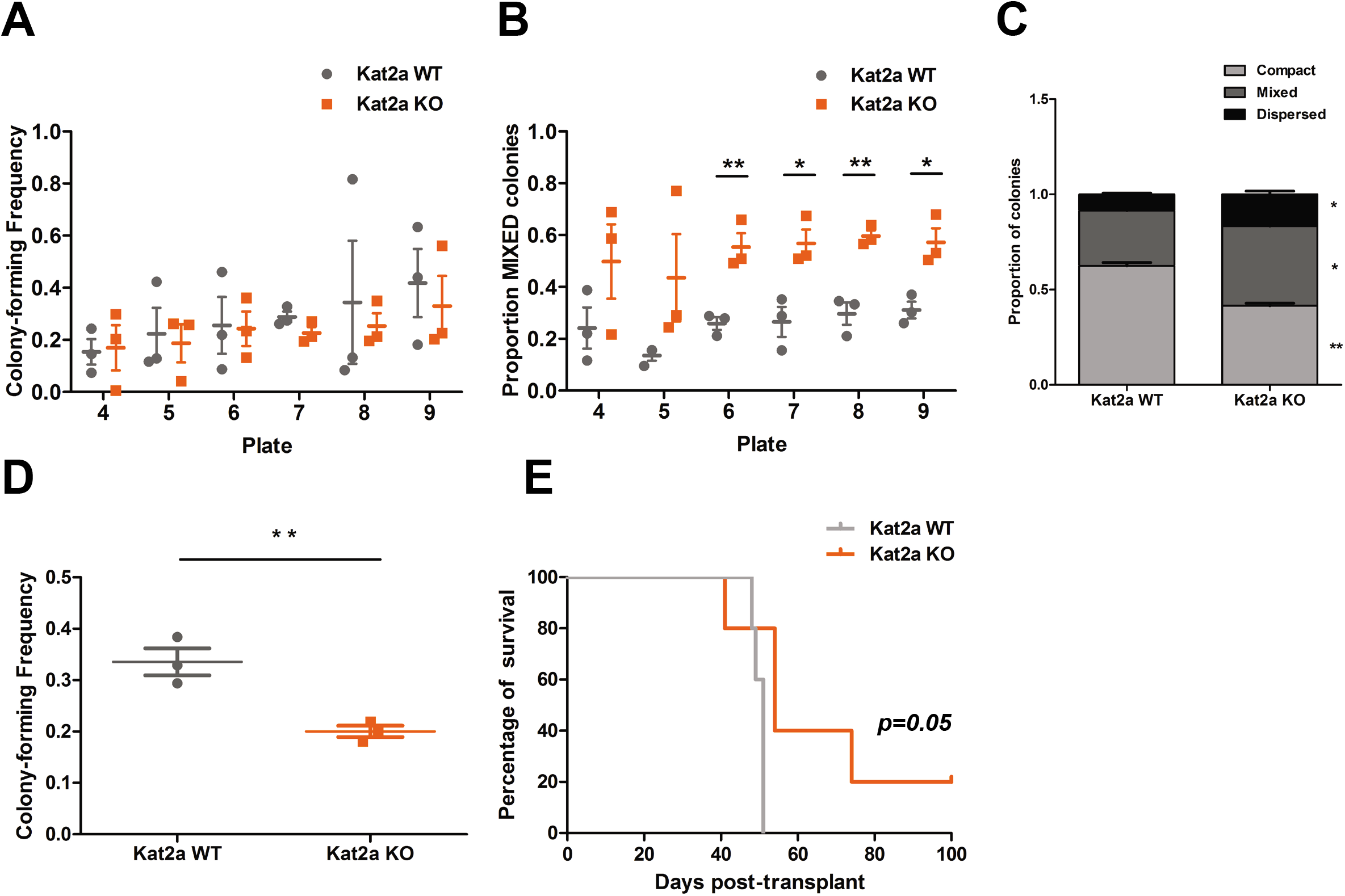
*Kat2a* depletion impairs establishment of MLL-AF9-initiated leukemia. **(A)** Serial replating CFC assays from in vitro MLL-AF9 transformed cells on WT and KO background, mean ± SEM, n= 3. **(B)** Proportion of Mixed type colonies in MLL-AF9 transformed cells with *Kat2a* WT or KO background, mean ± SEM, n= 3, * p<0.01 and ** p<0.001, 2-tailed t test. **(C)** Proportion of colony types distribution in CFC assays from *in vitro* transformed MLL-AF9 BM cell lines, mean ± SEM, n= 2-3, * p<0.01 and ** p<0.001 **(D)** CFC assay frequency from clonal liquid cultures of MLL-AF9 *in vitro* transformed, mean ± SEM, n= 3, ** p<0.001. **(E)** Survival curve of animals transplanted with MLL-AF9 transformed WT or KO cells. Log rank test was performed, mean ± SEM, n=5, p=0.05. Two-tailed t-test was performed in (B), (C) and (D).

We probed the effect of *Kat2a* in *MLL-AF9*-driven transformation *in vivo* by injecting lethally-irradiated recipients with WT and KO Lin-BM cells transduced for 2 days with retrovirus encoding the *MLL-AF9* oncogenic fusion. Animals developed leukemia 3 months after transplantation, as previously described, with a modest survival advantage for recipients of KO cells (Fig. 2E). At the point of culling, no differences in leukemia burden (Supplementary Fig. 2B), hematological parameters (Supplementary Fig. 2C) or in bone marrow cellularity (Supplementary Fig. 2D-E) were observed between genotypes. Taken together, the results suggest that Kat2a facilitates AML initiation, likely through inhibition of leukemia cell differentiation or promotion of self-renewal, and we sought to explore the molecular mechanisms behind this effect.

### Kat2a is required for promoter acetylation of general transcriptional and post-transcriptional regulators

Kat2a is a histone acetyl-transferase required for deposition of the activating H3K9 acetylation (ac) mark (Jin et al., 2014), and capable of catalysing multiple acetyl-modifications (Kuo and Andrews, 2013) at promoters and at enhancers (Krebs et al., 2011). Using chromatin immunoprecipitation followed by next-generation sequencing (ChIP-seq), we identified promoters as H3K4 tri-methyl (me3) peaks (Supplementary Fig. 3A) and enhancers with H3K4 mono-methyl (me1)-enriched regions (Supplementary Fig. 3B) and inspected the pattern of distribution of H3K9ac in promoter and enhancer elements in MLL-AF9 primary leukemia initiated by *Kat2a* KO or WT cells. Although global H3K9ac was minimally changed between genotypes, there was a specific depletion of H3K9ac peaks at promoters in regions devoid of concomitant H3K27ac activation mark (Fig. 3A). Conversely, H3K9ac was mildly increased at candidate active enhancer regions marked by the presence of H3K27ac (Fig. 3B), suggesting a possible pattern of imbalance of H3K9ac regulation between promoters and enhancers.

**Figure 3.**
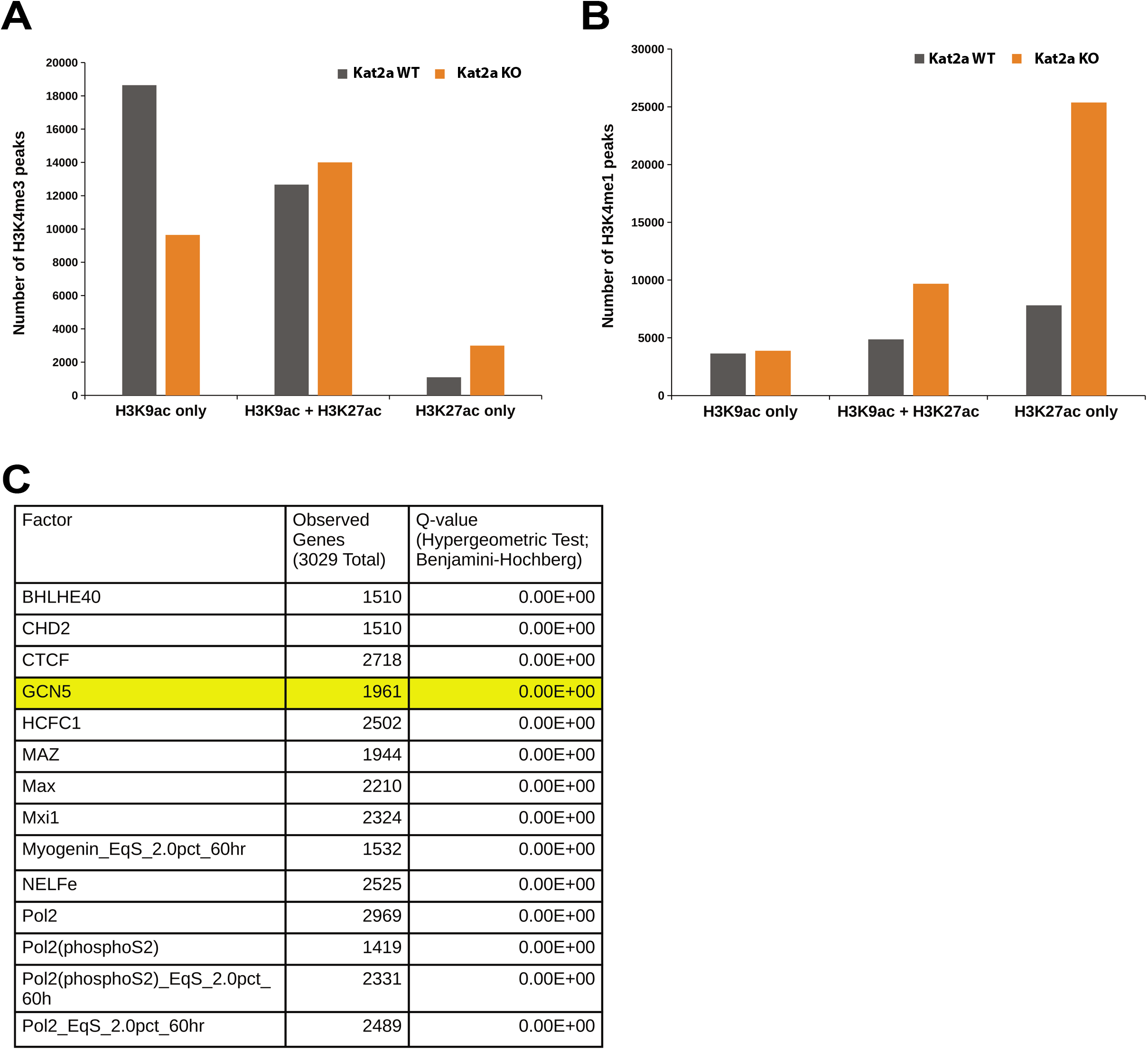
Loss of *Kat2a* depletes H3K9ac in a subset of promoters. **(A)** Distribution of H3K9ac peaks at promoters independently and in association with H3K27ac peaks. **(B)** Distribution of H3K9ac peaks at enhancers independently and in association with H3K27ac peaks. **(C)** ENCODE enriched DNA binding factors for genes associated with differentially-acetylated promoter peaks. List represents factors with Q-value equal to 0, ordered by proportion of observed H3K9ac differential genes to total number of binding site-associated genes. Long list is provided in Supplementary File 1.

We focused on those promoter peaks with unique loss of H3K9ac upon *Kat2a* depletion, and used the ENCODE database (Auerbach et al., 2013) to confirm enriched experimental binding of KAT2A (aka GCN5) in other model systems (Fig. 3C and Supplementary File 1). Similar to a previous study of the effects of *Kat2a* and H3K9ac loss in embryoid bodies (Wang et al., 2018), we also found evidence for increased representation of MYC targets, which is a known Kat2a interacting protein (Hirsch et al., 2015). Genes associated with differentially-acetylated promoter peaks fell into 3 main categories (i) general transcription factor and transcription initiation machinery; (ii) RNA metabolism, splicing and translation; and (iii) mitochondrial metabolism (Supplementary File 2). Of note, a quarter of the direct MLL-AF9 targets (Bernt et al., 2011) are differentially H3K9 acetylated upon *Kat2a* loss (Supplementary Fig. 3C and Supplementary File 3), including some (*Hoxa7*), but not all (*Hoxa9, Hoxa10* and *Meis1*) of the redundant Homeobox leukemia self-renewal gene signature. This raises the possibility that Kat2a may exert its effects through general, rather leukemia-specific, transcriptional and post-transcriptional regulatory mechanisms.

### Differential H3K9ac subsequent to Kat2a loss results in transcriptional variability

The role of yeast Gcn5 as a regulator of locus-specific intrinsic transcriptional noise suggested that loss of *Kat2a* might also increase transcriptional noise at H3K9ac target loci, an idea supported by our recent observations in mouse ES cells upon Kat2a inhibition (Moris et al., 2018). We approached this problem by single-cell transcriptional profiling of WT and KO primary leukemia cells on a high-throughput 10X Genomics platform. We pooled samples from 4-5 primary leukemias of each genotype, sorted live GFP+ cells reporting the presence of the *MLL-AF9* fusion, and successfully sequenced over 4000 cells *Kat2a* WT and KO cells each, for a total of 13166 transcripts. Basic measures of gene alignment and quality control are summarized in Supplementary Fig. 4A. In order to minimize the confounding effect of rarely or very low expression genes, we filtered out transcripts detected in less than 10% of all cells, and only included cells where a minimum of 500 different transcripts were detected, thus focusing the analysis on what we considered a ‘Robust’ gene set. These were a total of 2588 genes (Supplementary File 4), sampled from 7360 cells (3835 WT, 3525 KO). The Robust gene set captured the functional categories enriched amongst differential H3K9ac targets (Supplementary File 5), with significant overlap of individual genes (Supplementary File 6), indicating the validity of integrated analysis.

Differential gene expression analysis between WT and KO cells identified genotype-specific distributions within the Robust gene set, which corresponded to minimum changes in average gene expression. These were nevertheless predominantly of down-regulation upon *Kat2a* excision, as might be anticipated from loss of a histone acetyl-transferase (Fig. 4A). We observed a significant increase in gene expression variability as measured by coefficient of variation (CV=standard deviation/mean), which was apparent at all levels of gene expression (Supplementary Fig. 4B). Despite aggregate similarity of average gene expression between WT and KO AML cells, we sought to correct for its putative influence on the CV differences, by using distance to the median (DM) as an alternative variability measure (Kolodziejczyk et al., 2015). Notably, genes with promoter H3K9ac dependent on the presence of Kat2a and experimental evidence of Kat2a binding on chromatin as per the ENCODE database (henceforth called ‘Kat2a targets’), displayed a unique gain in transcriptional variability (Fig. 4B), that was not observed amongst non-targets after correction for average gene expression. Interestingly, the Kat2a-dependent variable genes exhibited lower basal variability than non-Kat2a targets, suggesting that Kat2a may function to maintain stable levels of gene expression. The general regulatory nature of the acetylation target genes is compatible with this view. In addition, motif analysis of Kat2a-dependent promoters revealed the enriched presence of a long unknown A/T rich motif (Fig. 4C), which we also found upon re-examination of H3K9ac-depleted promoters in mouse ES cells with Kat2a inhibition (data not shown). It is possible that this motif represents a nucleosome displacement region (Dadiani et al., 2013), thus accounting for the minimally variable and robust expression (Supplementary Fig. 4C) from these promoters, such as previously reported for some of the represented gene categories including ribosomal proteins (Field et al., 2008; Tirosh and Barkai, 2008).

**Figure 4.**
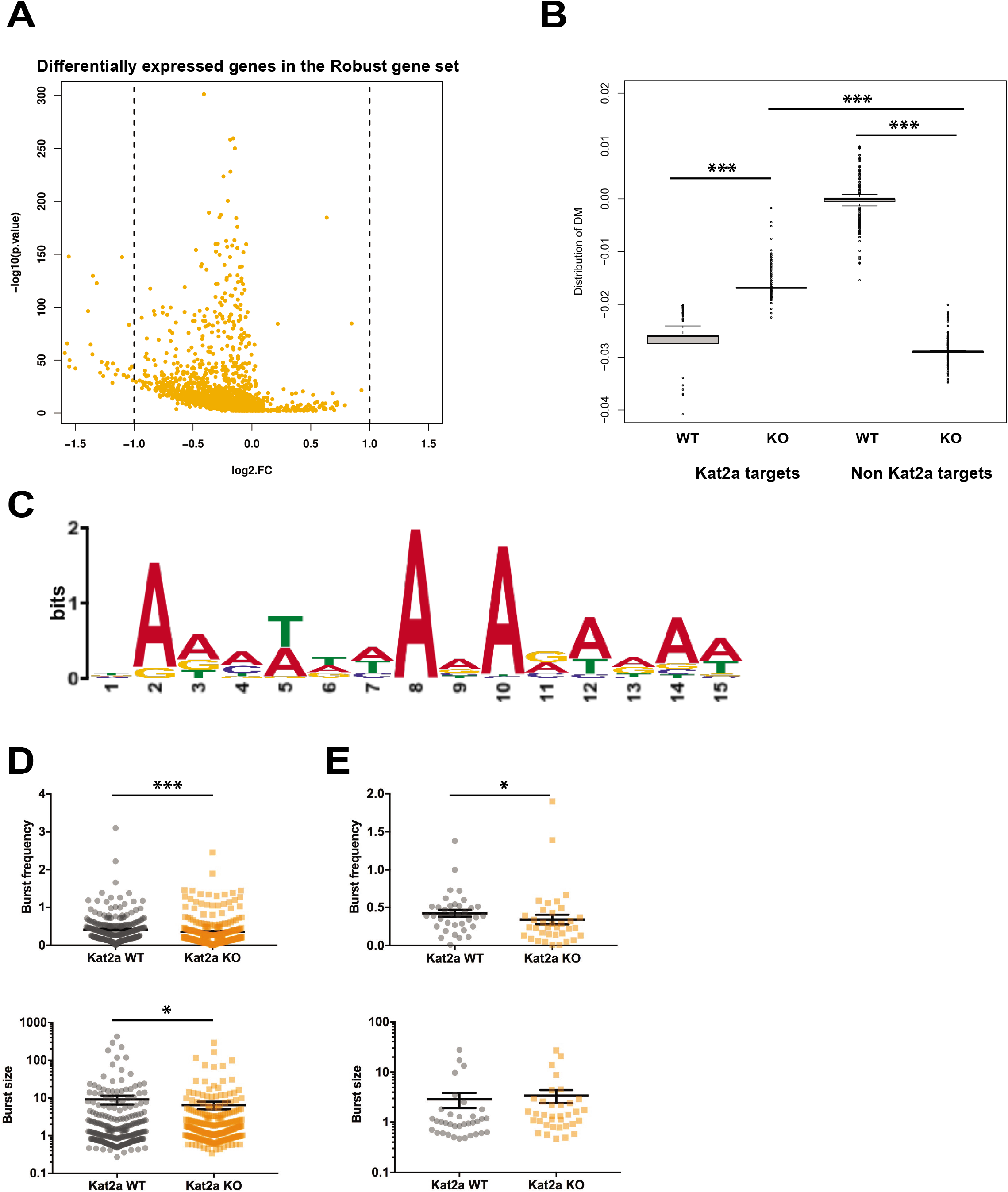
*Kat2a* KO leukemia cells have increased transcriptional heterogeneity. **(A)** Volcano plot of differentially expressed genes between *Kat2a* WT and KO primary leukemic cells as analysed by 10X Genomics single cell RNAseq. DE genes (Butler et al., 2018) in colour. **(B)** Distance to the median measure of transcriptional heterogeneity in *Kat2a* WT and KO single leukemic cells. Kat2a target and non-target genes analysed separately, with bootstrapping. The comparison was performed using non-parametric Kolmogorov-Smirnov test to compare distance to the median between *Kat2a* WT and *Kat2a* KO genotypes at 0.05 level of significance. All comparisons significant p<10^−5^. **(C)** MEME enriched motif in Kat2a target gene promoters. **(D)** Estimated burst frequency (top) and burst size (bottom) parameters for genes in the Robust gene set in *Kat2a* WT and KO primary leukemic cells. Parameters derived by applying D3E algorithm to single cell RNAseq data. **(E)** Estimated burst frequency (top) and burst size (bottom) parameters for Kat2a target genes. In (D) and (E), *p<0.05, ***p<0.001, computed with Wilcoxon rank sum test with continuity correction.

### Kat2a regulates transcriptional bursting activity in cells with stem-like characteristics

Having established that loss of *Kat2a* associates with increased cell-to-cell variability in expression levels of a subset of directly-targeted genes, we asked whether the variability reflected differential regulation of locus transcriptional bursting, and hence modulation of transcriptional noise. We made use of the D3E code developed by the Hemberg lab (Delmans and Hemberg, 2016) to parameterize and model differential gene expression from single-cell RNA-seq data (Supplementary Fig. 4D). Model fitting captured the contribution of burst frequency, but not burst size, to gene expression CV, whilst both parameters contributed significantly to mean gene expression (Supplementary Fig. 4E), supporting the validity of the analysis. Parameter fitting to the Robust gene set revealed significant decreases in both size and frequency of bursting in KO cells (Fig. 4D), with consequent reduction of average gene expression and gain in CV (Supplementary Fig. 4F). Notably, when focusing on Kat2a targets, we observed a pattern of significantly reduced frequency of bursting and a trend towards a gain in burst size (Fig. 4E), suggesting that Kat2a loss uniquely affects burst frequency and may achieve the observed gain in cell-to-cell variability through control of transcriptional noise. Indeed, the CV of Kat2a targets was increased, but we did not observe changes in average Kat2a target gene expression (Supplementary Fig. 4G). In an attempt to validate the results on a more homogenous subset of cells, and avoid confounding effects of cellular heterogeneity, we used dimensionality reduction and clustering strategies to compartmentalize WT and KO AML cells into clusters with identical transcriptional composition. We applied the RACE-ID algorithm (Grun et al., 2016) to the combined total of 7360 WT and KO cells, and optimally separated them into 12 clusters on the basis of 500 most highly variable genes in each genotype (Fig. 5A). The 2 genotypes displayed differential cluster-association patterns, with clusters 7, 11 and 12 being relatively underrepresented amongst *Kat2a* KO cells (respectively 0.5, 0.2 and 0.5 of *Kat2a* WT) (Supplementary Fig. 5A). We interrogated the individual clusters in terms of information entropy and inter-cluster connectivity as a means of predicting the stem-like or differentiated nature of their respective gene expression programs. Specifically, the STEM-ID algorithm (Grun et al., 2016), which builds on RACE-ID, defines a ‘stemness’ score on the assumption that stem cells exhibit a multitude of incipient lineage-affiliated programs (high information entropy), which are shared (high connectivity) with more differentiated cells. Cluster 7 had the highest ‘stemness’ score (Fig. 5B and Supplementary Fig. 5B), and we focused on it for model fitting. Consistent with the global findings, modelling of transcriptional parameters for Kat2a targets using the cells in cluster 7 revealed significantly lower frequency of bursting and associated high CV in KO cells (Fig. 5C and Supplementary Fig. 5C). Again, we observed a mild gain in burst size (Fig. 5C), which associates with unchanged mean expression levels (Supplementary Fig. 5C). In contrast, modelling of cells in cluster 6, with the lowest STEM-ID score, revealed no differences in transcriptional parameters between WT and KO cells (Fig. 5D). Of note, Kat2a targets had lower average gene expression in cluster 6 (Supplementary Fig. 5D). Overall, the data suggest that Kat2a target genes associate with candidate stem-like clusters and that Kat2a regulates their expression through buffering of transcriptional variability.

**Figure 5.**
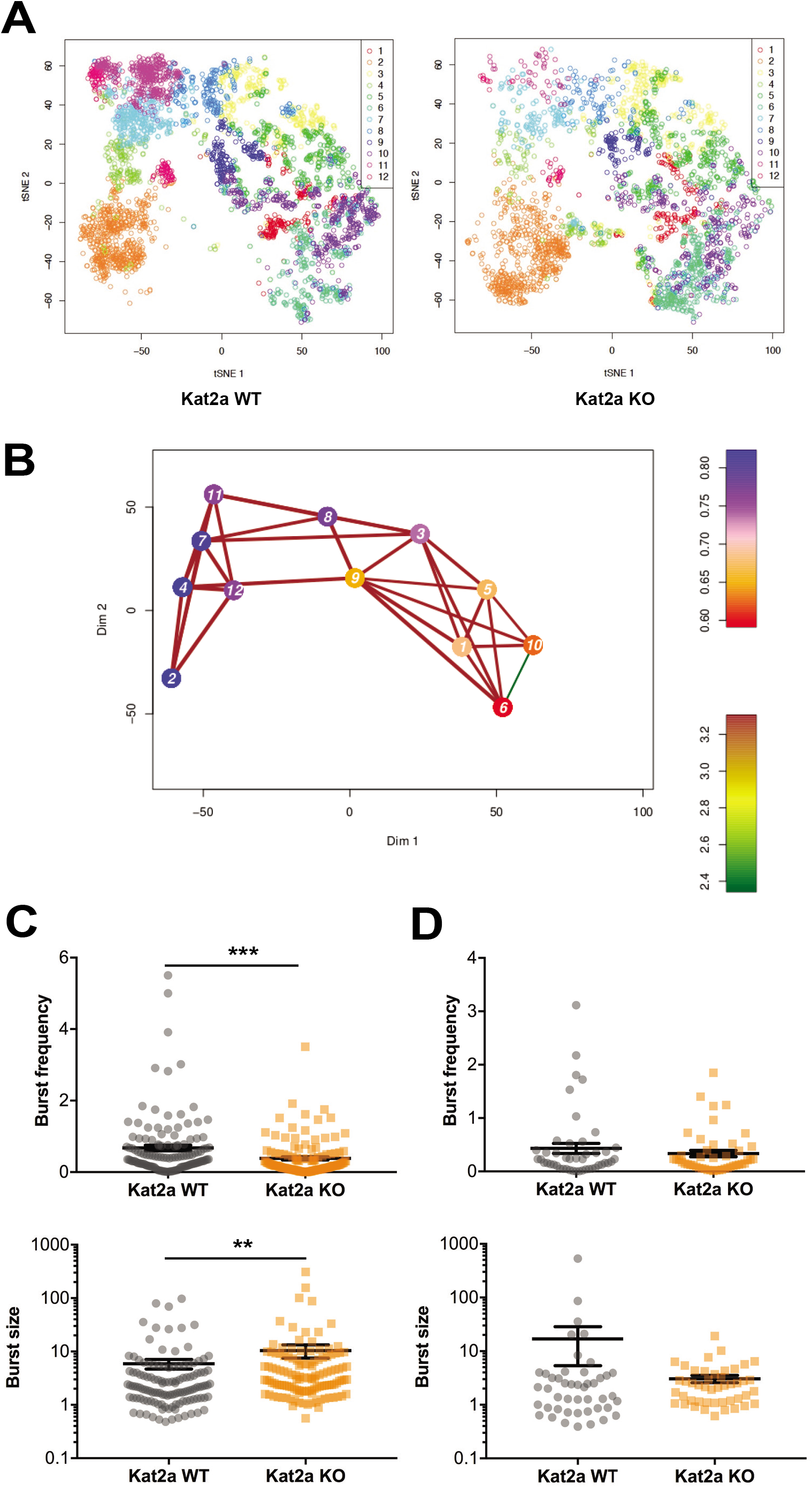
Transcriptional architecture of *Kat2a* KO AML suggests depletion of compartments with stem-like characteristics. **(A)** t-SNE plot of single cell RNA-seq data combining *Kat2a* WT (left) and KO (right) leukemic cells using the most highly variable genes from each genotype. RACE-ID based clustering is color-coded with cells of each genotype displayed separately. **(B)** STEM-ID trajectory plot of analysis in A representing combined measures of information entropy and cluster connectivity strength; clusters as in A**. (C)** and **(D)** Burst frequency and size parameters for Kat2a targets in clusters 7 **(C)** and 6 **(D)** in panel A. Parameterization as per Fig. 4D-E; p-value<0.001 in both cases computed with Wilcoxon rank sum test with continuity correction.

### Kat2a regulates the activity of translation-associated genes

Having established that Kat2a loss results in deregulation of transcriptional activity with decrease of bursting frequency, we asked if this effect was biased towards particular classes of genes. Indeed, we found that translation-associated genes, including ribosomal protein genes and translation initiation factors, were significantly overrepresented (Fig. 6A) amongst Kat2a target genes in cluster 7 which responded with increased noise to *Kat2a* KO. Also, this translation-associated gene-signature was more highly expressed amongst cells in high STEM-ID clusters (Fig. 6B), suggesting a role in maintenance or propagation of the leukemia self-renewal state. In order to determine putative consequences of this transcriptional deregulation, we inhibited KAT2A activity in the *MLL-AF9* cell line MOLM13, and observed a dramatic reduction in polysome abundance, and a minor decrease in the 60S subunit (Fig. 6C), the latter putatively matching biased but not exclusive targeting of large subunit ribosomal protein genes. Since generalized targeting of ribosomal protein genes by *Kat2a* loss precludes the design of a classical single-gene phenotyping rescue experiment, we attempted to phenocopy aspects of *Kat2a* depletion in *MLL-AF9* transformed cells by inhibiting the translation machinery with the S6K1 inhibitor PF4708671, which targets the TOR pathway and impedes translation initiation and elongation. Treatment of *MLL-AF9* transformed primary mouse BM cultured cells with PF4708671 resulted in a relative enrichment of mixed and disperse colonies in colony-forming assays (Fig. 6D), as observed upon *in vitro* transformation of *Kat2a* KO cells. Total colony formation from transformed primary cell lines was reduced upon S6K1 inhibition, which could also be seen in lines established from *Kat2a* KO cells (Fig. 6E).

**Figure 6.**
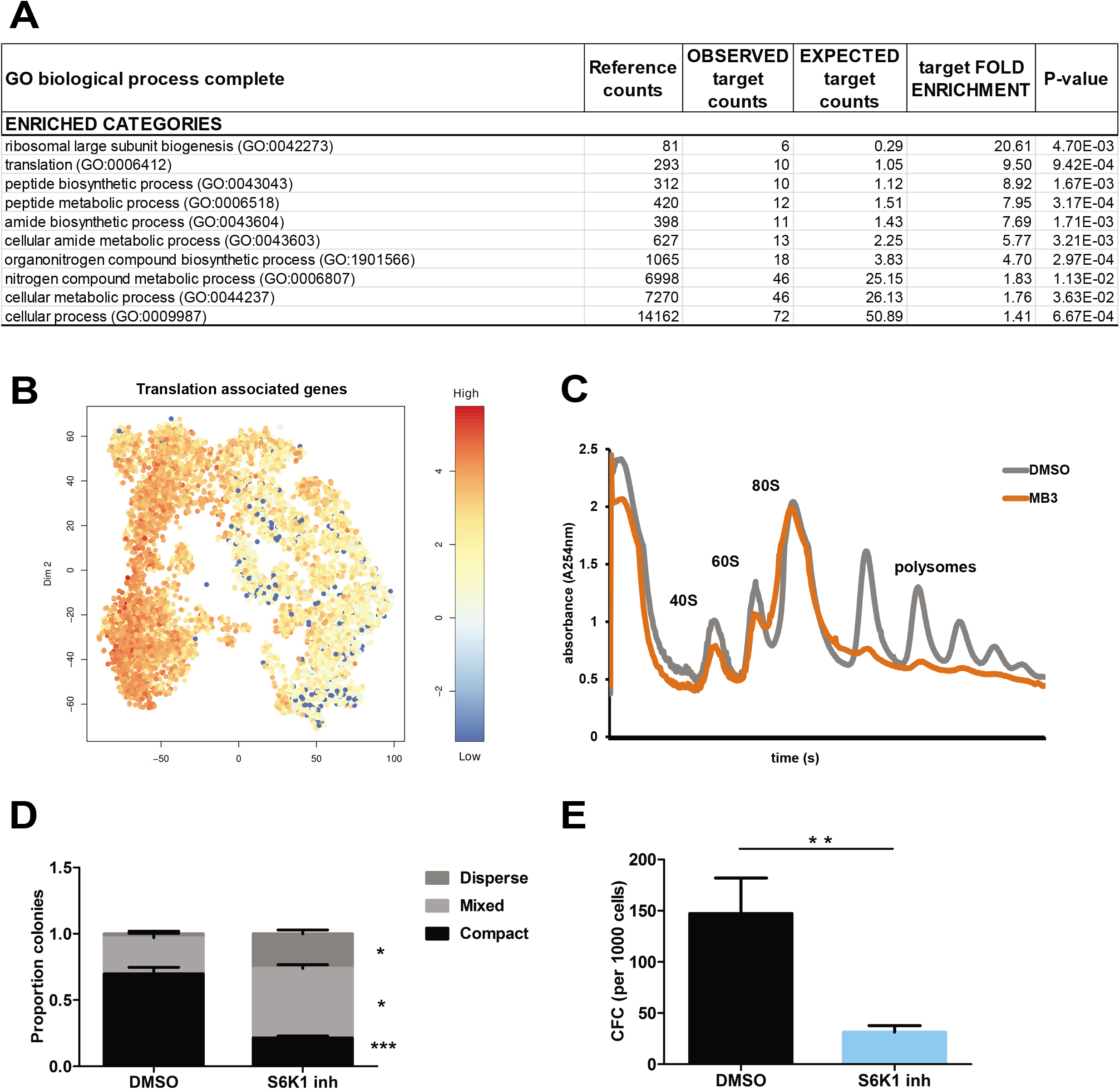
Translation-associated programs contribute to the *Kat2a*-regulated AML phenotype. **(A)** GO biological process enriched categories in *Kat2a* target genes with low burst frequency (cluster 7). Significant enrichments used Binomial analysis with Bonferroni correction in PANTHER (Mi et al., 2017). **(B)** Heatmap representation of the translation associated signature with reduced burst frequency in *Kat2a* KO cells (cluster 7). tSNE plot of single-cell expression as in Fig. 5A. **(C)** Polysomal profiling of MOLM-13 cells (carrying the *MLL-AF9* fusion gene) upon overnight treatment with DMSO or the Kat2a inhibitor MB-3 (200uM, (Biel et al., 2004)); data are representative of 2 independent experiments. **(D)** Proportion of colonies in *Kat2a* WT cells treated with DMSO and S6K1 inhibitor (mean + SEM, n=3, *** p< 0.001, *p<0.05, paired t-test). **(E)** Colony forming efficiency in *Kat2a* WT cells treated with DMSO and S6K1 inhibitor (mean + SEM, n=3, ** p< 0.01, paired t-test).

### Kat2a loss depletes functionalMLL-AF9 leukemia stem-like cells

Finally, we asked whether the enhanced transcriptional variability observed in STEM-ID high cluster 7 upon *Kat2a* loss, not only resulted in molecular drift of KO cells along candidate differentiation trajectories, but also corresponded to functional loss of leukemia stem-like cells. For this, we quantified the frequency of functional stem cells in primary *Kat2a* WT vs. KO MLL-AF9 leukemia through *in vivo* limiting dilution analysis. Specifically, we transplanted cell doses of 500 to 50000 live GFP+ cells obtained from the pooled primary leukemia samples of each genotype into secondary recipients and observed a survival advantage in animals transplanted with *Kat2a* KO cells, which was significant at the lowest 500-cell dose (Fig. 7A). The estimated frequency of leukemia stem-like cells (LSC) was significantly reduced in *Kat2a* KO-initiated MLL-AF9 leukemia: 1:17363 (95% confidence interval, 95CI: 4913-63312; vs. WT=1:714 (95CI: 183-2791) (Fig. 7B). Unlike our observations in primary leukemia, flow cytometry surface marker analysis of secondary leukemia samples indicated an overall gain of differentiated monocytic Mac1+Gr1-cells (p<0.01), but not granulocytic Mac1+Gr1+ cells, consistent with the lineage affiliation of MLL-AF9 leukemia (Fig. 7C). Accordingly, there was a trend towards depletion of Lineage-Kit+Sca1-CD34+FcgR+ leukemic GMP-like (L-GMP) cells (p=0.07) in *Kat2a* KO secondary MLL-AF9 leukemia (Fig. 7C). The fact that this was not significant may suggest a partial dissociation between LSC phenotype and function, or highlight the presence of LSC with alternative cell surface phenotypes.

**Figure 7.**
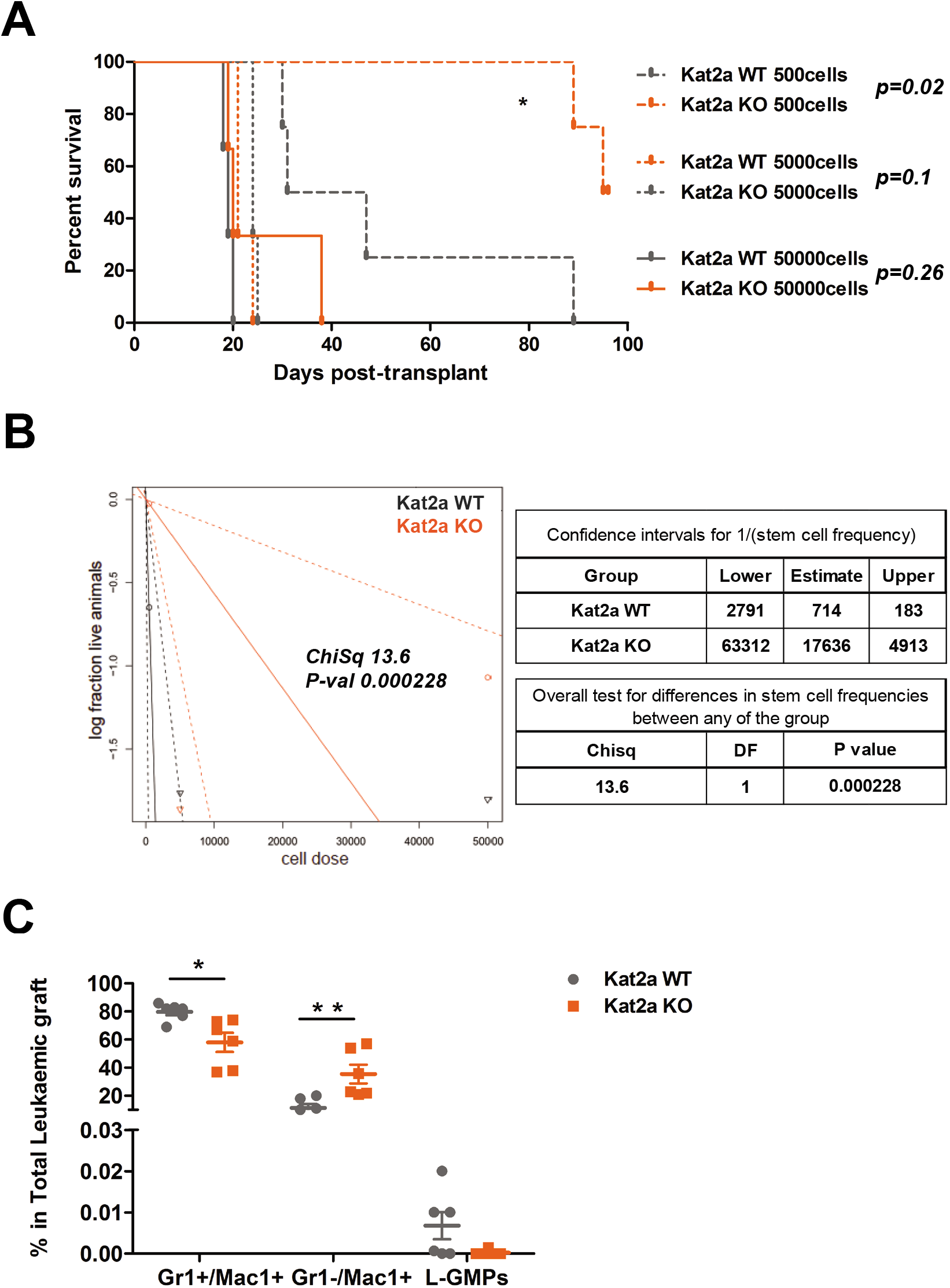
Kat2a loss depletes functional MLL-AF9 leukemia stem-like cells. **(A)** Survival curve of secondary recipients of MLL-AF9 leukemic cells from WT and KO backgrounds. Transplantation of limiting dilution cell doses as indicated. Log rank test for difference in survival, n=3-4/per dose group. 50000 cells p=0.26, 5000 cells p=0.1, 500 cells p=0.02. **(B)** Extreme Limiting Dilution Analysis (ELDA, (Hu and Smyth, 2009)) of leukemia-initiating cell frequency in *Kat2a* WT and KO primary leukemia. Data as in A. **(C)** Flow cytometry analysis of BM cells from secondary WT and KO leukemia transplant recipients (50000 and 5000 cells). L-GMPs are Gr1-Mac1-cKit+Sca1-CD34+FcgR+, mean ± SEM, n=6, Student’s t test was performed, ** p< 0.001, *p<0.01, and L-GMPs p=0.07.

## DISCUSSION

In this study, we link regulation of transcriptional variability with leukemia cell fate through knockout of histone acetyl-transferase Kat2a. Using an *MLL-AF9* model of mouse AML initiated through transplantation of retroviral-transduced BM progenitor cells, we show that *Kat2a*-depleted cells have impaired preservation or propagation of self-renewal, which dramatically reduces functional leukemia stem-like cell content of primary tumors without notable alterations in cell surface phenotype. Transformation *in vitro* proceeds despite a bias towards differentiation, but *Kat2a* KO transformed cell lines have decreased colony-initiating potential. Similarly, establishment of primary KO leukemia is only mildly delayed with little evidence of differentiation. This nevertheless becomes apparent upon secondary transplantation, concomitantly with the reduction in leukemia stem-like cell content. Interestingly, we note a dissociation between surface phenotype and stem-like function, suggesting that identification of the classical L-GMP surface antigen phenotype (Krivtsov et al., 2006) may not absolutely associate with function. Also, importantly, we, like others, did not observe that loss of *Kat2a* introduced changes to normal hematopoiesis (Bararia et al., 2016), particularly in HSC, LMPP or GMP compartments that directly contribute to *MLL-AF9* transformation. Repeated 5-FU treatment or secondary transplantation (data not shown) also failed to reveal a stem cell function defect, indicating specific dependency of leukemia stem-like cells on expression of Kat2a.

We used single-cell transcriptional analysis to capture cell-to-cell heterogeneities within seemingly phenotypic equivalent primary AML from both genotypes. Whilst differences between genotypes were minimal in terms of average gene expression, we identified a clear distinction in cell-to-cell variability in transcript levels that was specific to a subset of promoters characterized by H3K9, but not H3K27, acetylation, and which were dependent on Kat2a activity. Compatible with the proposed role of H3K9ac in stabilizing, but not initiating, transcription (Jin et al., 2014), the increase in transcriptional variability was not matched by a decrease in mean expression levels; moreover the respective genes were expressed at least at equivalent average levels to the remaining loci analyzed by single-cell RNA-sequencing irrespective of genotype. Given the absence of H3K27ac at these promoters, our analysis may have identified a group of promoters that does not have enhancer capacity in alternative states or cell contexts, a hypothesis worth exploring in future work. Compatible with this notion, we observed that, unlike the unique Kat2a-dependent promoter subset, promoters marked by H3K9ac and H3K27ac were relatively enriched in *Kat2a* KO leukemia cells. The same was observed at H3K4me1-positive enhancers. One hypothesis is that the Kat2a-dependent promoters with exclusive H3K9ac are bound by Kat2a in the context of SAGA, whilst promoter and enhancer regions with dual H3K9ac and H3K27ac regions are bound by ATAC. ATAC is a distinct Kat2a-containing complex previously associated with promoters not bound by SAGA as well as with enhancers, and that possesses a second acetyl-transferase activity (Krebs et al., 2011). It is also possible that loss of *Kat2a* may be partially compensated by Kat2b (aka, Pcaf) activity, but with locus selectivity. We did not find evidence of enhanced Pcaf expression in *Kat2a* KO leukemia (data not shown), but that does not exclude that it can have enhanced activity.

It is well established that most, if not all, eukaryotic genes have a discontinuous mode of transcriptional activity, described as transcriptional bursting. Briefly, transcriptional bursting depends on the kinetics of locus activation and inactivation, and of RNA production and degradation, with the number of mRNA molecules produced during a ‘burst’, or activity cycle, determining the gene expression level, and the frequency of activation of gene promoters influencing the breadth of transcript levels present in a cell through time, in other words transcriptional noise, or snapshot heterogeneity. In the case of our data, modelling of promoter status and transcriptional bursting activity suggested that the regulatory differences observed between *Kat2a* KO and WT cells were specifically due to a loss in the frequency of bursting activity, which correlates with transcriptional noise. The number of mRNA molecules produced by each burst, or burst size, on the other hand, was not changed or was even mildly increased upon loss of *KAT2A*, suggesting a mechanistic link between H3K9ac and bursting frequency. This recapitulates findings in yeast linking H3K9ac at gene promoters with noise, but not level of gene expression (Weinberger et al., 2012), and provides mechanistic insight into the cell-to-cell heterogeneity elicited by *Kat2a* loss. In a recent study, the Naëf lab has shown that locus-specific manipulation of promoter, but not distal or enhancer, H3K27 acetylation can change transcriptional bursting frequency (Nicolas et al., 2018). Whilst the association with H3K27ac is unclear in our study, there is a clear contribution of H3K9 acetylation to bursting frequency, which matches an association of promoter H3K9ac, in addition to H3K27ac, and frequency of locus activation in the Naëf study (Nicolas et al., 2018). The mild gain in burst size, although unproductive in terms of transcriptional level, could reflect the differential reconfiguration of H3K9ac at promoters and enhancers upon *Kat2a* loss, and will be interesting to follow-up in subsequent studies. Indeed, our lab has recently developed a KAT2A-Cas9 fusion capable of catalyzing targeted acetylation events (data not shown), that will be instrumental in answering these questions.

In linking the promoter-specific effects of Kat2a on H3K9ac and frequency of transcriptional bursting to the observed depletion of leukemia stem-like cells we found that general metabolic categories, in particular related to RNA processing, rather than known leukemia-associated programs, were affected in their chromatin signature and frequency of bursting. While we cannot exclude that our focus on loss, rather than reduction, of H3K9ac, combined with current limitations of single-cell RNA sequencing in capturing low-expressed genes, may have missed individual candidates, the agreement between the two levels of analysis, and indeed the ontology overlap with published studies of Kat2a-depleted or inhibited ES cells (Hirsch et al., 2015; Wang et al., 2018), including ours (Moris et al., 2018), suggest that Kat2a may regulate pervasive, rather than cell specific programs. The identification of a candidate nucleosome displacement motif in Kat2a target promoters also indicates specific regulation of highly and widely expressed genes. Amongst these, we found that translation as a category was targeted by *Kat2a* depletion, and demonstrated that not only is the assembly of polysomes perturbed by Kat2a inhibition, but that perturbation of the translational machinery can re-capture defects in *in vitro* propagation of leukemia-initiating cells akin to those imposed by *Kat2a* depletion. In agreement, Morrison and collaborators (Signer et al., 2014) have reported that impaired protein synthesis upon genetic depletion of the ribosomal protein machinery impedes leukemia self-renewal, whilst having non-linear dose-dependent effects on normal hematopoiesis, mimicking our own observations in the *Kat2a* KO setting. Future studies directing Kat2a histone acetylation activity to single or multiple loci will illuminate individual *vs*. global target gene contributions to the leukemia phenotype. However, it is tempting to speculate that the generic nature of the programs impacted by Kat2a at the level of transcriptional noise may configure an underlying propensity towards execution of cell fate transitions, which can be of a different nature in different biological contexts. Analysis of the impact of Kat2a target programs in other malignant and normal stem cell systems, or at different stages of leukemia progression will test this hypothesis. It will also be interesting to determine if other candidate regulators of transcriptional noise, including pharmacological agents (Dar et al., 2014), produce similar effects and can be exploited therapeutically in AML, as well as in other hematological and non-hematological malignancies.

## EXPERIMENTAL PROCEDURES

### Tables of key reagents

Table A – Primers used for genotyping and analysis of *Kat2a* excision

Table B – Antibodies used in flow cytometry analysis and cell sorting

Table C – Flow cytometry gating strategies

Table D – Taqman assays for quantitative RT-PCR analysis of gene expression

Table E – Chromatin immunoprecipitation buffers and solutions

Table F – Antibodies used in chromatin immunoprecipitation

### Generation and analysis of *Kat2a* conditional knockout mice

*Kat2a^F1/F1^* conditional knockout mice (Lin et al., 2007) were bred with *Mx1-Cre*^+/−^ transgenic mice (Kuhn et al., 1995), in a C57Bl/6 background. Littermates were genotyped for *Kat2a* LoxP sites and for *Mx1-Cre:* ear notch biopsies were digested using KAPA express extract (Sigma Aldrich) and KAPA2G ROBUST HS RM Master Mix (2x) (Sigma Aldrich). PCR cycling protocol: 95C, 3min; 40x (95°C, 15sec; 60°C, 15sec; 72°C, 60sec); 72°C, 60sec. Primers listed in [Table A]. DNA products were run on a 1% Agarose Gel in TAE (1x), at 100V and visualized using an AlphaImager UV transilluminator (Protein Simple). Cre-mediated recombination was induced in 6-10-week-old mice by administration of 5 alternate-day intraperitoneal injections of poly(I)-poly(C) (pIpC), 300μg/dose. After pIpC treatment, animals were identified as *Kat2a* WT = *Kat2a^F1/F1*^ Mx1-Cre^−/−^* and *Kat2a* KO = *Kat2a^F1/F1*^ Mx1-Cre ^+/−^*. Excision efficiency was determined by qPCR of genomic DNA (gDNA) from Peripheral Blood (PB), Spleen (Sp) or Bone Marrow (BM). gDNA was extracted using Blood and Tissue DNA easy Kit (Qiagen) and quantified by Nanodrop (Thermo Scientific). qPCR analysis used Sybr Green Master Mix (Applied Biosystems) and two sets of primers (Table A, Fig. 1A): *Kat2a*-IN, a/b, within the excised region; *Kat2a*-OUT, c/d distal to the second LoxP site. Expression levels were determined by the Pfaffl method following normalization to *Kat2a*-OUT. PB was collected by saphenous vein and differential blood cells counts were determined using a Vet abc automated counter (Scil Animal Care, Viernheim, Germany). Mice were kept in an SPF animal facility, and all experimental work was carried out under UK Home Office regulations. Animal research was regulated under the Animals (Scientific Procedures) Act 1986 Amendment Regulations 2012 following ethical review by the University of Cambridge Animal Welfare and Ethical Review Body (AWERB).

**Table A.**
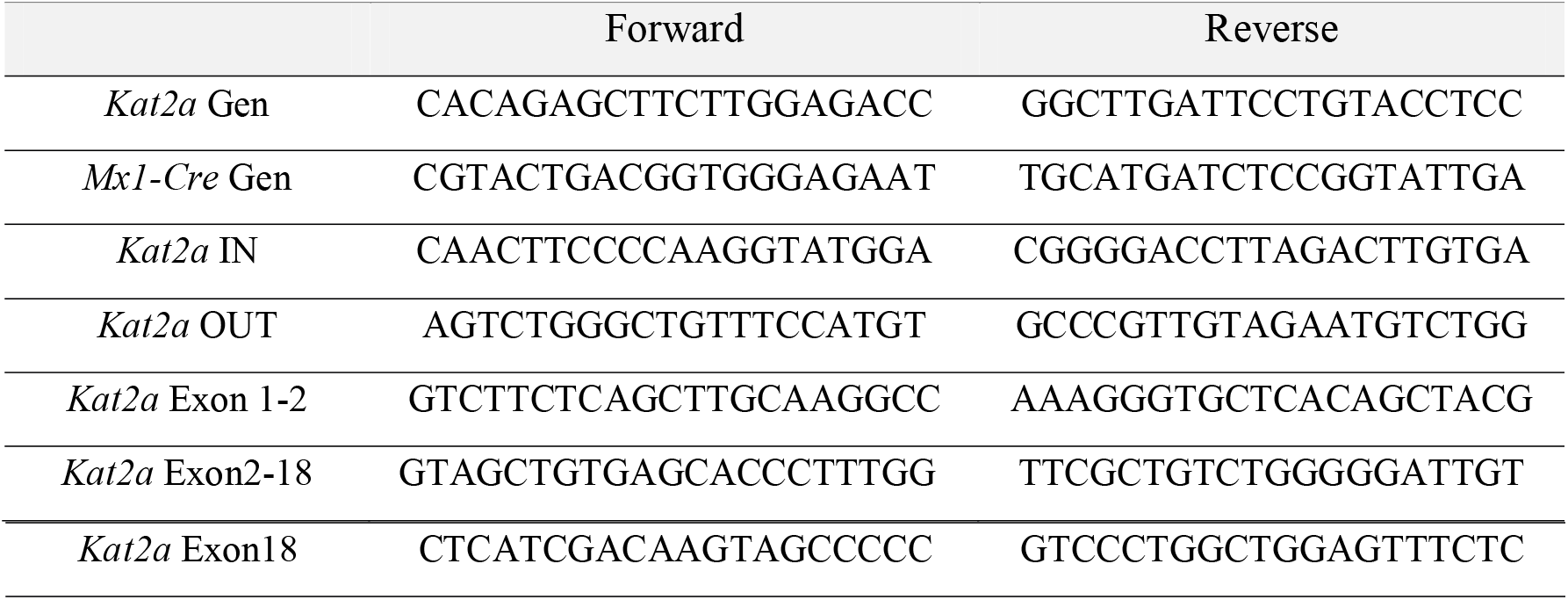
Primers used for genotyping and analysis of *Kat2a* excision

### Isolation of mouse BM stem and progenitor cells

BM was isolated from mouse long bones as described before (Pina et al., 2015). Following red blood cell lysis, total BM suspension was depleted of differentiated cells using a cocktail of biotinylated lineage antibodies (Table B) and streptavidin-labeled magnetic nanobeads (Biolegend), according to manufacturers’ instructions. Cells were directly used in transplants, colony-forming assays or flow cytometry for analysis of normal hematopoiesis. For leukemia studies, cells were cultured overnight at 37°C 5% CO2 in RPMI supplemented with 20% Hi-FBS (R20), 2mg/mL L-Glutamine, 1% PSA, 10 □ng/mL of murine Interleukin 3 (mIL3), 10 □ng/mL of murine Interleukin 6 (mIL6), and 20 □ng/mL of murine Stem Cell Factor (mSCF) (cytokines from Peprotech) (supplemented R20), followed by retroviral transduction.

**Table B.** Antibodies used in flow cytometry analysis and cell sorting

| Antibody | Fluorochrome | Catalogue # | Clone | Dilution | Supplier |
| --- | --- | --- | --- | --- | --- |
| CD45.1 | FITC | 110705 | A20 | 1:200 | Biolegend |
| CD45.2 | AF700 | 56-0454-81 | 104 | 1:200 | Ebioscience |
| B220/CD45R | Biotin | 116204 | RA3-6B2 | 1:300 | Biolegend |
| TER119 | Biotin | 103204 | Ter119 | 1:300 | Biolegend |
| Gr1 | Biotin | 108404 | RB6-8C5 | 1:300 | Biolegend |
| CD3e | Biotin | 100304 | 145-2C11 | 1:300 | Biolegend |
| CD11b | Biotin | 101204 | M1/70 | 1:300 | Biolegend |
| CD11b/Mac1 | PE-Cy7 | 25-0112-81 | M1/70 | 1:200 | Ebioscience |
| FcγR | PE | 101308 | 93 | 1:100 | Biolegend |
| CD34 | APC | 128612 | HM34 | 1:100 | Biolegend |
| cKIT/CD117 | APC-Cy7 | 105826 | 2B8 | 1:50 | Biolegend |
| Sca1 | PE-Cy7 | 108114 | D7 | 1:100 | Biolegend |
| Gr1 | PB | 108430 | RB6-8C5 | 1:100 | Biolegend |
| Flt3/CD135 | PE | 135305 | A2F10 | 1:100 | Biolegend |
| CD105 | PE | 562759 | MJ7/18 | 1:100 | BD |
| CD150 | Af647 | 562647 | Q38-480 | 1:100 | BD |
| CD41 | Biotin | 13-0411-81 | eBioMWReg30 | 1:100 | Ebioscience |
| Streptavidin | BV405 |  |  | 1:200 | Biolegend |

### Colony forming cell (CFC) assays

For analysis of normal progenitors, sorted mouse BM cells were plated at a density of 200-400 cells/plate in duplicates, in MethoCult GF M3434 (STEMCELL Technologies). Colonies were scored at 7-9 days. For analysis of MLL-AF9 leukemia, retroviral-transduced BM cells were plated in M3434 at an initial density of 10000 cells/condition and scored and re-plated every 6-7 days. Re-plating was performed up to passage 9, with 4000 cells/condition used from plate 3. CFC assays from mouse MLL-AF9 transformed lines were seeded in M3434 and scored 6-7 days later. RPS6K inhibition studies were set by adding 3.3uL DMSO, either as vehicle or with a final concentration of 3.5uM of PF4708671 (Tocris), directly to the methylcellulose medium, with mixing prior to cell addition.

### *In vivo* analysis of leukemia initiation and engraftment

For analysis of normal hematopoiesis, 10^6^ *Kat2a* WT or *Kat2a* KO cKit+ cells were intravenously injected via tail vein into lethally irradiated (2*5.5Gy) CD45.1 recipient mice. At the described time-points, BM and Sp were collected and processed into a single-cell suspension for surface marker staining and flow cytometry analysis. For AML induction, we transplanted 1.5×10^6^ cKit+ *Kat2a* WT or *Kat2a* KO cells transduced with *MSCV-MLL-AF9-IRES-YFP*, intra-venous into lethally irradiated (2* 5.5Gy) CD45.1 recipient mice. Upon signs of illness and following human end-point criteria, animals were culled, tissue samples collected for histology analysis, and BM and Sp processed into single-cell suspensions. Flow Cytometry analysis and DNA extraction were performed. For limiting-dilution analysis, 5*10^2^ - 5×10^4^ cells from primary leukemia pooled BM of each genotype were transplanted into sub-lethally irradiated (1*5.5Gy) CD45.1 recipient mice (3-4/dose and genotype).

### Retroviral transduction

Retroviral construct *MSCV-MLL-AF9-IRES-YFP* was previously described (Fong et al., 2015). For viral particle production, Human Embryonic Kidney (HEK) 293T cells were seeded at 2.5×10^6^ cells/10cm dish in DMEM supplemented with 10% Hi-FBS, 2mg/mL L-Glutamine, 1% PSA and cultured overnight at 37°C 5% CO2. The following day, a transfection mix [per plate: 47.5 uL of TranSIT (Miros), 5ug of packaging plasmid psi Eco vector (5ug), retroviral vector (5ug) and 600uL of Optimem Medium (Gibco)] was prepared according to manufacturer’s instructions and added dropwise to cells followed by plate swirling and overnight culture at 37°C 5% CO2. Medium was replaced with R20 the next day. At 24 and 48 hours after R20 replacement, medium was collected and filtered through a 0.45uM syringe filter, and viral particle suspension medium was added to BM cells. BM cells from 6-10 week-old *Kat2a* WT and *Kat2a* KO mice were collected and Lineage-depleted as described above (Isolation of mouse BM stem and progenitor cells), and cultured overnight at 37°C 5% CO2 in supplemented R20. For viral transduction, BM cells were briefly centrifuged at 400G, 5 minutes, and viral particle suspension medium supplemented with 10 □ng/mL mIL3, 10 □ng/mL mIL6, and 20 □ng/mL mSCF added to a final density of 10^6^ cells/mL. Cells were plated in 6-multiwell plates and centrifuged for 1 hour at 2000rpm, 32°C. After, cells were incubated for 4 hours at 37°C 5% CO2. A second round of viral transduction was performed, with post-centrifugation incubation performed overnight. Next day, cells were collected, pelleted and washed three times with PBS (2x) and R20 (1x). YFP level was accessed by Flow Cytometry in a Gallios Analyser (Beckman Coulter).

### Establishment of MLL-AF9 transformed cell lines

MLL-AF9 clonal liquid cultures were set up using *MLL-AF9* retrovirus-transduced primary BM cells (see Retroviral Transduction section). Transformed cells enriched *in vitro* by 3 rounds of serial plating (CFC assays) were maintained in R20 supplemented on alternate days with mSCF, mIL3 and mIL6, all at 20ng/mL. Cells were cultured at 2*10^5^ cells/ml and passaged when they reached a density of 1*10^6^/ml.

### Flow Cytometry

Cell surface analysis of BM and Sp was performed using a panel of antibodies described on Table B according to sorting strategies detailed on Table C. Data was acquired using a Gallios Analyser (Beckman Coulter) and analysis performed in Kaluza software (Beckman Coulter). For sorting, an Influx or an AriaII BD sorter were used.

**Table C.** Flow cytometry gating strategies

| Cell population | Sorting Strategy |
| --- | --- |
| HSC | Lin<sup>-</sup> cKit<sup>+</sup> Sca1<sup>+</sup> CD34<sup>-</sup> Flt3<sup>-</sup> |
| MPP | Lin<sup>-</sup> cKit<sup>+</sup> Sca1<sup>+</sup> CD34<sup>+</sup> Flt3<sup>-</sup> |
| LMPP | Lin<sup>-</sup> cKit<sup>+</sup> Sca1<sup>+</sup> CD34<sup>+</sup> Flt3<sup>+</sup> |
| CMP | Lin<sup>-</sup> cKit<sup>+</sup> Sca1<sup>-</sup> CD34<sup>+/low</sup> CD16/32<sup>low</sup> |
| GMP | Lin<sup>-</sup> cKit<sup>+</sup> Sca1<sup>+</sup> CD34<sup>+</sup> CD16/32<sup>high</sup> |
| MEP | Lin<sup>-</sup> cKit<sup>+</sup> Sca1<sup>+</sup> CD34<sup>-</sup> CD16/32<sup>-</sup> |
| Lin Negative | CD3e<sup>-</sup> B220<sup>-</sup> Gr1<sup>-</sup> CD11b<sup>-</sup> Ter119<sup>-</sup> |

### Quantitative Real time PCR (Q-RT-PCR)

Total RNA was extracted using Trizol Reagent (Invitrogen). RNA from equal numbers of cells was reverse-transcribed using Superscript II (Invitrogen), following manufactures’ instructions. Complementary (c)DNA was analyzed in duplicate or triplicate by qPCR using Taqman gene expression assays (Table D) and Taqman Gene Expression Mastermix (Applied Biosystems). Gene expression levels were determined by the Pfaffl method following normalization to Reference gene, as stated. For exon 2-18 in-frame products, qPCR using Sybr Green Master Mix (Applied Biosystems) was performed in triplicates.

**Table D.** Taqman assays for quantitative RT-PCR analysis of gene expression

| Gene Symbol | Assay ID |
| --- | --- |
| *Ppia* | Mm03024003_g1 |
| *Hprt* | Mm01545399_m1 |
| *Kat2a* | Mm00517402_m1 |

### Polysomal profiling

MOLM-13 cells were grown to an approximate density of 1 × 10^6^ cells/mL, treated with cycloheximide (100 μg/mL) for 15 minutes, washed in ice-cold PBS and stored at −80°C. Cells were lysed in buffer A (20mM HEPES pH 7.5, 50mM KCl, 10mM (CH_3_COO)2Mg, EDTA-free protease inhibitors (Roche), supplemented with cycloheximide 100 μg/mL, 1mM PMSF, 100 U/mL RNase inhibitor (Promega), 1% (vol/vol) sodium deoxycholate, and 0.4% (vol/vol) NP-40) at 10^8^ cells/mL for 10 minutes on ice. Lysates were cleared by centrifugation (8000g for 5 minutes at 4°C) and 3 A254nm units loaded onto a 10%-50% (wt/vol) sucrose gradient in buffer A in Polyallomer 14 × 95 mm centrifuge tubes (Beckman). After centrifugation (Beckman SW40Ti rotor) at 260 900g for 3 hours at 4°C, gradients were fractionated at 4°C using a Gilson Minipulse 3 peristaltic pump with continuous monitoring (A254nm) and polysome profiles recorded using a Gilson N2 data recorder.

### Chromatin Immunoprecipitation sequencing (ChlP-seq)

Total BM cells from duplicate pools of MLL-AF9 *Kat2a* WT and *Kat2a* KO primary leukemia samples were crosslinked with 1% Formaldehyde Solution (Sigma Aldrich) for 10 min at room temperature (RT), with gentle rotation (50rpm). Fixation was stopped with Glycine, and cells incubated for 5 min, RT, with gentle rotation (50rpm), followed by two washing steps in ice-cold PBS. Cell pellets were resuspended in Lysis buffer (Table E) followed by Nuclei preparation. Chromatin pellets were sheared in a Bioruptor Pico Plus (Diagenode) in TPX tubes, using 3 runs of 11 cycles (Cycle: 30sec ON 30sec OFF) on high setting. A short spin was performed between runs and samples were transferred to new TPX tubes. 1:10 of total sheared chromatin was kept for input reference. Immunoprecipitation was set up using Dilution Buffer, Protease cocktail Inhibitor, and the respective antibody (Table F) and the sheared chromatin incubated overnight at 4°C with rotation. On the following day, protein A/G magnetics beads were pre-cleared with Dilution Buffer supplemented with 0.15% of SDS and 0.1%BSA, then mixed with immunoprecipitation mix and incubated for at least 4hours at 4°C with rotation. Chromatin-Antibody-Beads mixes were sequentially washed with ChIP Wash1, ChIP Wash2, ChIP Wash3 (Table E) and captured on a magnetic rack. Captured beads were incubated for 20 minutes with rotation in freshly prepared Elution Buffer. Supernatants were collected and decrosslinking performed overnight. DNA was column-purified using DNA clean and concentrator TM 5 KIT (Zymo Research), according to manufacturer’s instructions, using 20uL Zymo Elution Buffer. DNA quality control was performed using DNA Qubit 2.0/3.0 (Invitrogen). Libraries were prepared at the MRC/WT Cambridge Stem Cell Institute Genomics Core Facility using NextFlex Rapid DNA kit 12 cycles, and sequenced on a HiSeq 4000 sequencer at the CRUK Cambridge Research Institute.

**Table E.**
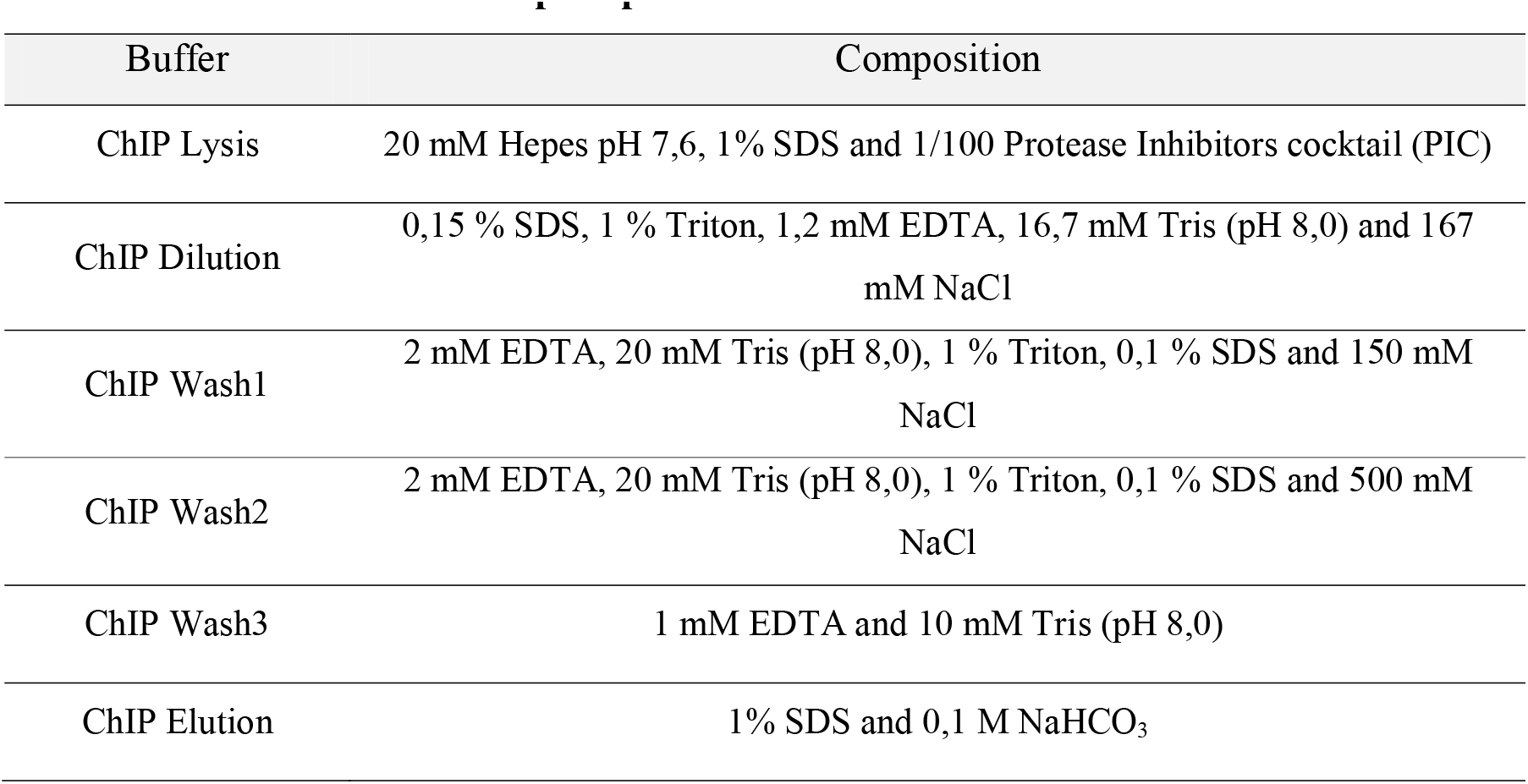
Chromatin immunoprecipitation buffers and solutions

**Table F.** Antibodies used in chromatin immunoprecipitation

| Antibody | Catalogue# | Supplier |
| --- | --- | --- |
| H3K27ac | Ab4729 | Abcam |
| H3K9ac | 07-352 | Millipore |
| H3K4me3 | Ab8580 | Abcam |
| H3K4me1 | Ab8895 | Abcam |

### ChIP-seq data analysis

Four histone modification marks, specifically H3K4 tri-methylation (me3), H3K4 mono-methylation (me1), H3K27 acetylation (ac) and H3K9ac were studied in *Kat2a* WT and *Kat2a* KO samples using ChIPseq analysis. The experiments were performed in duplicate. Raw ChIPseq reads were analyzed on the Cancer Genomics Cloud (CGC) platform (Lau et al., 2017). Reads were aligned to the mouse mm10 Genome obtained from UCSC genome browser using the Burrows-Wheeler Aligner (BWA). Peaks from the aligned reads were obtained using the MACS2 peak calling algorithm with a significance q-value of 0.05. The deepTools bamCoverage command (Ramirez et al., 2016) was used to compare the enrichment of reads in the ChIPseq samples relative to corresponding controls. ChIP-seq samples with distinct separation between control and sample pair for a given marker were retained with exclusion of one H3K4me1 and one H3K27ac replicate. To analyze the changes in acetylation patterns at promoter and enhancer elements, H3K4me3 and H3K4me1 peaks from WT and KO were crossed with H3K9ac-only peaks, H3K27ac-only peaks and dual H3K9ac and H3K27ac peaks from the corresponding genotypes. The H3K9ac-only peaks associated with me3 (promoter elements) were used for further analysis. Genomic peaks were obtained for *Kat2a* WT and *Kat2a* KO genotypes separately using Bedtools intersect (Quinlan and Hall, 2010) and H3K4me3 K9ac peaks exclusive to WT genotypes retained as putative Kat2a peaks. Peak locations were converted to fastq sequences using UCSC table browser tool (Karolchik et al., 2004). Genomic Regions Enrichment of Annotations Tool (McLean et al., 2010) was used to assign gene identities to the fastq sequences associated with putative Kat2a peaks. Using the GREAT tool, the genomic region for gene identification was restricted to 1kb upstream and 500bp downstream of the transcription start site (TSS) to infer genes regulated at the promoter level. We used ENCODE ChIP-Seq Significance Tool (Auerbach et al., 2013) to obtain putative transcription factors regulating these targets, as well as lists of genes experimentally bound by GCN5/KAT2A, to confirm the identity of putative Kat2a targets. MEME-chip tool version 4.12.0 (Bailey et al., 2009) was used to define DNA motifs associated with these genes. This gene list was considered to represent putative Kat2a targets.

### Single-cell RNA sequencing

Terminal BM samples from *Kat2a* WT and *Kat2a* KO MLL-AF9 primary leukemia animals were collected (WT - 5; KO - 4) and the individual cell samples stored at −150°C. Cells were thawed and pooled for library preparation. 12K live cells per genotype pool were sorted on an Influx sorter (BD) on the basis of YFP expression (reporting *MLL-AF9*), Hoechst 32258 exclusion (live cells) and singlet configuration (pulse width) and used for library preparation aiming at 6K single cells per sample. The libraries were prepared using 10X Genomics platform using single-end sequencing on a NextSeq 500 sequencer at CRUK Cambridge Research Institute. Raw single cell RNAseq fastq reads (sequenced on 10X Genomics platform) were analysed using Cellranger software (v2.1.1) to obtain the cell-gene count-matrix. Seurat (Butler et al., 2018) was used for pre-processing the count-matrix data and obtaining differential gene expression between the two genotypes. RaceID/StemID (Grun et al., 2016) algorithms were used for clustering using t-SNE and obtaining pseudo-temporal arrangement of clusters based on entropy information and cluster stem scores. Parameters for the stochastic gene expression were fitted to the two-state promoter model using the D3E algorithm (Delmans and Hemberg, 2016). R scripts were written for plotting the results as boxplots and for bootstrapping of distance-to-median measure between *Kat2a* WT and KO.

### Statistical analysis

Statistical tests performed are specified in the figure legends. Differences were obtained at p-value significant at 0.05. All analyses were performed in statistical language R (version 3.4.4).

### Data deposition

All single-cell RNAseq data and ChIPseq data were deposited in GEO (SuperSeries GSE118769).

## AUTHOR CONTRIBUTIONS

Conceptualization: C.P.; Methodology: A.F.D., R.K., G.G., B.J.H., S.P., C.P.; Formal analysis: R.K., S.P.; Investigation: A.F.D., R.K., S.G., ST., E.F., R.R.A., K.Z.; Resources: C.P., S.P., B.J.H.; Writing – Original Draft: C.P.; Writing – Review & Editing: C.P., B.J.H., S.P., G.G., S.G.; Visualization: A.F.D., R.K., S.G.; Funding Acquisition: C.P.; Supervision: C.P.

## ACKNOWLEDGEMENTS

The *Kat2a (Gcn5) Flox* allele mouse strain was a kind gift from Prof Sharon Dent, M.D. Anderson Cancer Centre, Smithville, TX, USA. The authors are grateful to Dr Matt Wayland for help with Cell Ranger installation and initial processing of the 10X Genomics data; to the Flow Cytometry Facility at the Cambridge Institute for Medical Research, Cambridge for expert assistance with cell sorting; to the Genomics Core Facility at the MRC/WT Cambridge Stem Cell Institute for ChIP-Seq library preparation; to the CRUK Cambridge Research Institute Genomics Core Facility for 10X Genomics library preparation and Next-Generation Sequencing services; and to the Cambridge Biomedical Services for animal husbandry. This work was funded by a Kay Kendall Leukaemia Fund Intermediate Fellowship (KKL888) and by a Leuka John Goldman Fellowship for Future Science (2017) to C.P.. S.P. is funded through a Cambridge-DBT Lectureship; R.K. was funded by an Isaac Newton Trust (INT) Research Grant and a Wellcome Trust ISSF/INT/University of Cambridge Joint Research Grant to C.P.; S.G. is funded by a Lady Tata Memorial Trust PhD Studentship, a Trinity Henry Barlow Trust Scholarship, and the Cambridge Trust; K.Z. received funding from AIRC (Italian Association for Cancer Research) and is the current recipient of a European Commission Horizon 2020 Marie Sklodowska Curie Post-Doctoral Fellowship.

